# Sonic stimulation, and low power microwave radiation can modulate bacterial virulence towards *Caenorhabditis elegans*

**DOI:** 10.1101/351924

**Authors:** Priya Patel, Hiteshi Patel, Dhara Vekariya, Chinmayi Joshi, Pooja Patel, Steven Muskal, Vijay Kothari

**Affiliations:** Institute of Science, Nirma University, India; Eidogen-Sertanty Inc., USA

**Keywords:** Sonic stimulation, ‘Om’, Microwave, Athermal effect, Virulence, Antimicrobial resistance

## Abstract

*Caenorhabditis elegans* worms infected with different pathogenic bacteria were subjected to sonic treatment to investigate whether such sound treatment can exert any therapeutic effect on the infected worms. Sonic therapy corresponding to 400 Hz, and the divine sound ‘om’ were found to confer protective effect on this nematode worm in face of bacterial infection, particularly that caused by *Serratia marcescens* or *Staphylococcus aureus*. The observed effect seemed to occur due to influence of sound on bacteria, and not on the worm. In addition to this, effect of microwave exposure on bacterial virulence was also investigated, wherein microwave exposure was found to reduce virulence of *S. aureus* towards *C. elegans*.

## Introduction

Infectious microorganisms have plagued mankind since ancient times, and caused numerous deaths throughout the history of humans on earth. Older civilizations with limited understanding of these *invisible* enemies tried to combat the burden of infections through use of a variety of herbal and herbomineral preparations. Later twentieth century saw the dawn of the antibiotic era. Discovery of antibiotics greatly helped the human race in their fight against the pathogenic microorganisms. However, owing to the development of antimicrobial resistance (AMR), and its promiscuous transfer among various microbial species, it has now become clear that antibiotics alone cannot hold the infectious microbes back, and in addition to the development of new antimicrobials, we need to look for novel non-antibiotic ways to combat microbial virulence, which may not replace the antibiotic use, but may emerge as useful alternative measures. Among such alternative/complementary ways to compromise bacterial virulence, one approach includes the use of non-invasive therapeutics such as electromagnetic radiation (EMR) (Elsheakh, 2017), acoustic waves (Bandara *et al*., 2014), plasma torch (Thiyagarajan, 2011), etc.

Different parts of the EMR spectrum such as X-rays and UV are known to have deleterious effects on almost all forms of life. Among them, biological effects of microwaves (MW) have received relatively lesser attention from the research community. Particularly the athermal (non-thermal) effect of MW has remained a matter of debate and controversy in the scientific as well as public domain (Singh and Kapoor, 2014). While thermal effects of MW are well-established, literature contains papers arguing in favour of the athermal effects, as well as, those indicating otherwise (Leszczynski *et al*., 2002; Regel and Achermann, 2011).

Among the most common abiotic factors interacting with life forms, one is sound, which can be classified into infrasound (<20 Hz), audible sound (20-20,000 Hz), and ultrasound (> 20 KHz) (Kadam *et al*., 2016) While ultrasonic frequencies are known to have deleterious effect on microbial cells leading to their lysis, not much research has been done on microbial response to sonic stimulation. Though few reports (Ying *et al*., 2009; Shaobin *et al*., 2010; Aggio *et al*., 2012; Gu *et al*., 2013; Kim, 2016; Liu *et al*., 2016; Murphy *et al*., 2016) have accumulated in literature describing the effect of external sound field on microbial cells, the precise mechanism by which acoustic vibrations affect microbial forms of life, yet remains to be elucidated.

Previously we have reported results of our *in vitro* experiments describing (a) effect of low power MW on microbial growth/metabolism (Kushwah *et al*., 2013), protein synthesis (Mishra *et al*., 2013), enzyme activity (Dholiya *et al*., 2012), exopolysaccharide production (Kushwah *et al*., 2013), toxin production (Ramanuj *et al*., 2015), quorum-sensing (QS) regulated pigment production (Chaudhari *et al*., 2014), etc.; and (b) effect of sonic stimulation on microbial growth (Sarvaiya and Kothari, 2015; Shah *et al*., 2016), antibiotic susceptibility (Sarvaiya and Kothari, 2017), and QS-regulated pigment production (Kothari *et al*., 2017; Kothari *et al*., 2018). Mutagenic effect of MW has also been previously described by us (Gosai *et al*., 2013; Kothari *et al*., 2014). Since QS (process of intercellular communication among bacteria) regulates expression of multiple genes in bacteria, including those associated with virulence, and some QS-regulated pigments (e.g. pyocyanin) are believed to be involved in killing of the eukaryotic hosts by bacterial pathogens, and our previous research indicated the potential of low power MW and sound waves to modulate bacterial QS *in vitro*; we undertook this study to investigate whether sonic/ MW treatment can modulate virulence of pathogenic bacteria, employing the nematode worm *Caenorhabditis elegans* as the model host. Pathogenic strains used in this study were *Chromobacterium violaceum*, *Pseudomonas aeruginosa*, *Serratia marcescens*, and *Staphylococcus aureus. C. violaceum* is considered as an emerging pathogen (Kothari *et al*., 2017), *P. aeruginosa* and *S. aureus* are among the most infamous notorious human-pathogenic bacteria, whose antibiotic-resistant varieties have been listed among the pathogens of high/critical importance by WHO (World Health Organization), and so is the case with the family Enterobacteriaceae (http://www.who.int/en/news-room/detail/27-02-2017-who-publishes-list-of-bacteria-for-which-new-antibiotics-are-urgently-needed) to which *S. marcescens* belongs.

Additionally, we also investigated effect of sonic stimulation on growth of *Prevotella melaninogenica* and *Biofidobacterium bifidum*. The former bacterium belongs to the family Prevotellaceae, whose lower abundance has been observed in patients suffering from neurological problems like autism (Qiao *et al*., 2018), and Parkinson’s disorder (PD) (Scheperjans *et al*., 2015), along with increased *Bifidobacteria* and Enterobacteriaceae abundance (Petrov *et al*., 2017). Hence perhaps approaches for modulating abundance of Prevotellaceae, Enterobacteriaceae, and other bacteria like *Bifidobacteria* and *Lactobacillus*, whose correlation with neurological disease/ well-being has been indicated in literature, may prove of some significance in management of conditions like PD. Among two such bacteria selected for this study, *B. bifidum* is a human gut inhabitant of gram-positive type, widely used as a probiotic strain with different perceived health benefits (Picard *et al*., 2005; Ku *et al*., 2016). Bifidogenic effect of plant products (e.g. prebiotics) or physical agents (e.g. EMR, sound waves) can also be of relevance, as higher abundance of these bacteria has been associated with ‘positive mood’ (Sarkar *et al*., 2016). *P. melaninogenica* (synonymous with *Bacteroides melaninogenicus)* is an anaerobic gram-negative bacterium, part of common flora present in mouth (Könönen, 1993).

## Materials and Methods

### Test organisms

Details of all the bacteria used in this study are given in Table-1. *C. elegans* was maintained on Nematode Growing Media (NGM; 3 g/L NaCl, 2.5 g/L Peptone, 1 M CaCl_2_, 1M MgSo4, 5mg/mL cholesterol, 1M phosphate buffer of pH 6, 17g/L agar agar) with *E. coli* OP50 (procured from LabTIE B.V., JR Rosmalen, the Netherlands) as feed. Worm population to be used for the *in vivo* assay was kept on NGM plates not seeded with *E. coli* OP50 for three days, before being challenged with test pathogen.

**Table 1.**
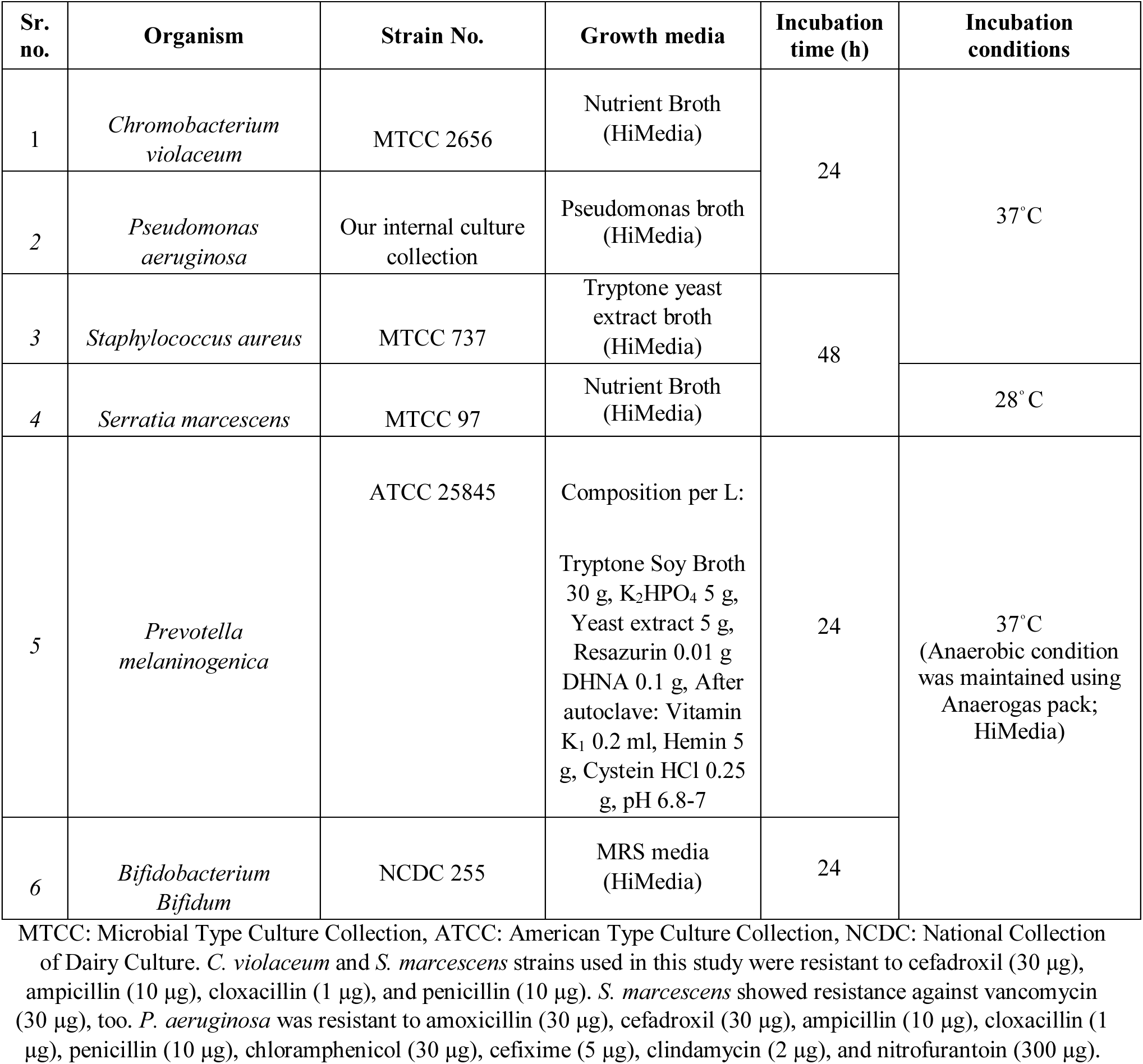
Test organisms

### Preparation of sound files

Two different sound types were used in this study. First type was containing the divine *Sanskrit* sound *‘Om’*, for which there is no translation. It is considered to be cosmic sound in Hindu culture (Gurjar and Ladhake, 2016). This sound file was sourced from (https://www.youtube.com/watch?v=yoYrLM5rGX8) which was then converted from mp4 to mp3 format using WavePad Sound Editor Masters Edition v 7.13. Total length of an individual ‘om’ sound in this file was 13.27 seconds. For the purpose of our experiment, we edited this file using WavePad Sound Editor Masters Edition v 7.13, so that a gap of 1 second occurs between two consecutive ‘om’ sound.

Another sound file contained the 400 Hz sound. This sound beep was generated using NCH^^®^^ tone generator. The sound file played during the experiment was prepared using WavePad Sound Editor Masters Edition v 7.13 in such a way that there is a time gap of one second between two consecutive beep sounds. Sound pertaining particularly to 400 Hz was selected, since in our previous experiments we found this frequency at 86 dB to be capable of modulating QS-regulated production of the pigment prodigiosin in *S. marcescens* (Kothari *et al*., 2016).

### Sonic stimulation

Inoculum of the test bacterium was prepared from its activated culture, in sterile normal saline. Optical density of this inoculum was adjusted to 0.08-0.10 at 625 nm (Agilent Technologies Cary 60 UV-vis spectrophotometer). This standardized inoculum (10%v/v) was used to inoculate suitable growth media (2 mL) for respective bacteria, followed by incubation under static condition at temp and for time length indicated in Table-1.

Following incubation density of the bacterial suspension was adjusted to OD_764_=1.50.One hundred μL of this bacterial suspension was mixed with 900 μL of M9 buffer (3 g/L KH_2_Po_4_, 6 g/L Na_2_HPo_4_, 5 g/L NaCl, 0.1% 1M MgSo_4_) containing 10 worms (L3-L4 stage). This experiment was performed in 24-well (sterile, non-treated) polystyrene plates (HiMedia TPG24), and incubation was carried out at 22 C. Two such 24-well plates were prepared, one to be treated as ‘control’, and another as ‘experimental’, wherein the former did not receive any sonic treatment, and the latter did. The ‘experimental’ plate was placed below a speaker (Minix sound bar, Maxtone Electronics Pvt. Ltd., Thane), with a distance of 13 cm between the speaker and the plate. This whole experimental set-up (Figure 1) was placed in a glass jar (Actira, L: 250 x W: 250 x H: 150 mm). Speaker was fitted between two side walls of the glass jar in such a way that its sound-emitting surface faces downside towards the 24-well plate lying on the floor of the jar. 24-well plates in both the glass jars were rotated by 180°once every day, so that all wells can be expected to receive almost equal sound exposure. Sound delivery from the speaker in experimental jar was provided throughout the period of incubation. Glass chambers were covered with a glass lid, and one layer of loose-fill shock absorber polystyrene, in such a way that the polystyrene layer gets placed below the glass lid. Silicone grease was applied on the periphery of the glass chamber coming in contact with the polystyrene material. This type of packaging was done to minimize any possible leakage of sound from inside of the chamber, and also to avoid any interference from external sound. Similar chamber was used to house the ‘control’ (i.e. not subjected to sound stimulation) plate. One non-playing speaker was also placed in the glass chamber used for the control plate at a distance of 13 cm from the plate, where no electricity was supplied and no sound was generated (Kothari *et al*., 2018). Intensity of sound was measured with a digital sound level meter (KUSAM-MECO^®^ KM 929 MK1; detection range 30-130 dB) by putting the sensor in place of the 24-well plate. The display on screen of this meter was visible through the glass of the experimental jars (which were sealed at the time of sound intensity measurement). Background noise (i.e. the sound produced by incubator) was also measured, whose value in the control and experimental jar was recorded to be 57.4 ± 0.23 dB and 58.8 ± 1.02 dB, respectively. When sound emission was on in the experimental chamber from speaker placed in it, dB values in the control chamber did not go beyond this background noise, indicating that sound from the experimental chamber did not reach the control chamber. Intensity of 400 Hz sound in the experimental chamber was set at 85±0.5 dB, and that of ‘om’ at 72-75 dB.

**Figure 1.**
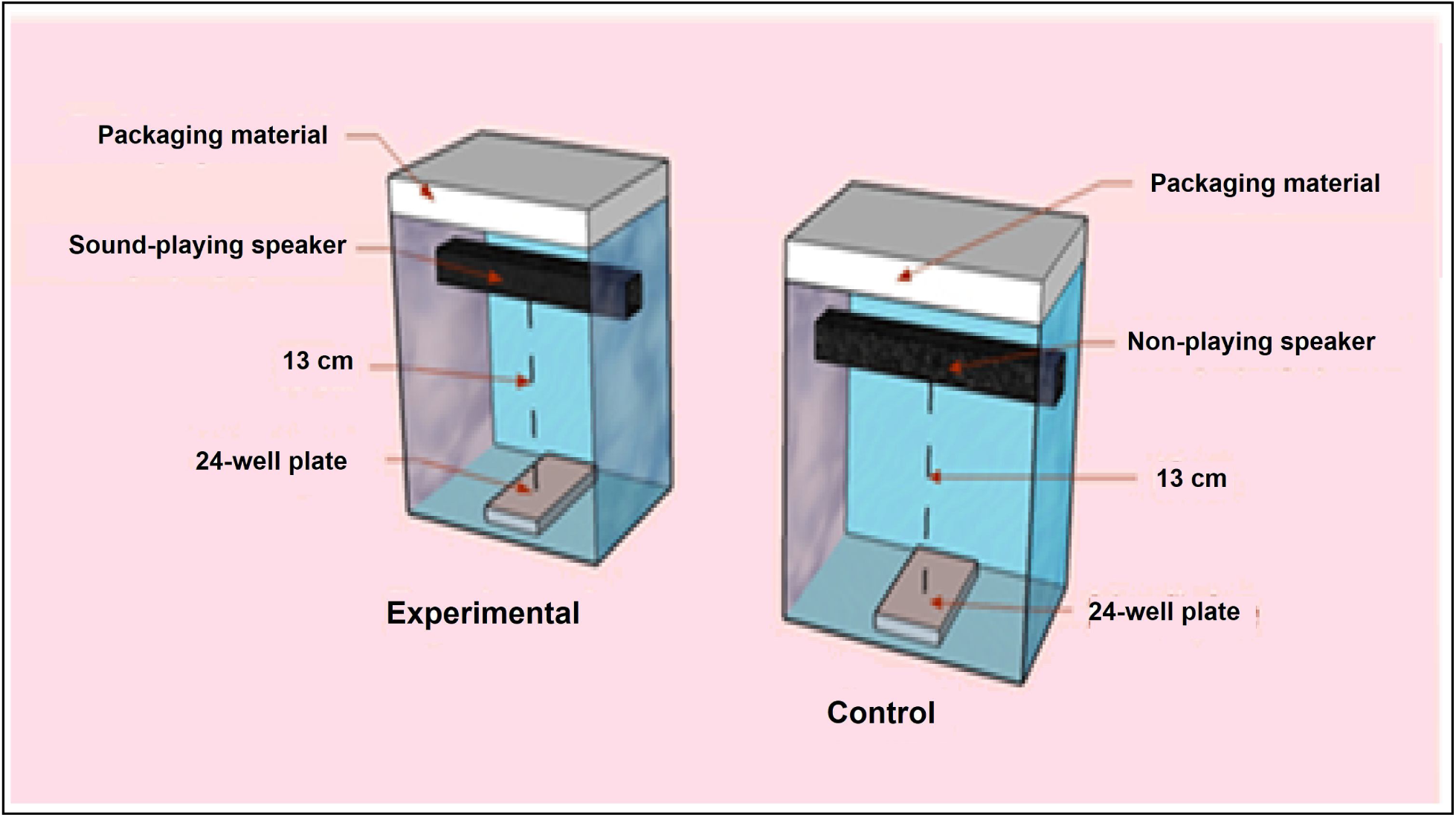
Experimental set up for investigating effect of sonic stimulation on infected *C. elegans*. Chambers were put in incubator taking care that neither they touch the incubator walls, nor to each-other, so that no vibrations are transferred.

Number of live vs. dead worms was counted everyday till five days by putting the plate (with lid) under light microscope (4X). Straight worms were considered to be dead. Plates were gently tapped to confirm lack of movement in the dead-looking worms. On last day of the experiment, when plates could be opened, their death was also confirmed by touching them with a straight wire, wherein no movement was taken as confirmation of death.

### Microwave (MW) treatment

Bacterial suspensions were prepared from an actively growing culture, in sterile normal saline, and its turbidity was adjusted to 0.08-1.00 at 625 nm. Test cultures (5 mL) in sterile screw capped glass vials (15 mL; Merck) were exposed to MW radiation (90 W; 2450 MHz) in a domestic MW apparatus (Electrolux^®^ EM30EC90SS) for 2, 4, or 6 min. Vials inside the MW apparatus were placed in a ice containing beaker (100 mL; Borosil^^®^^), so as to avoid any thermal heating. Temperature of the microbial suspension after MW treatment did not go beyond 15.6°C. (Table 2). Temp measurements before and after MW treatment were carried out using infrared thermometer (KUSAM-MECO^®^ IRL380). The whole MW treatment was performed in an air-conditioned room. Untreated inoculum was used as control (receiving no MW exposure, but ice treatment equivalent to that of experimental). Before MW treatment all the inoculum vials were put in ice for 5 min to nullify any variations in initial temperature. In case of control vial, put for 5 min in ice and then for 5 min at room temp (instead of MW treatment), the temp of inoculum was found to be 18°C. Test organisms were immediately (in less than 5 min) inoculated (at 10%v/v) into the growth medium following MW treatment, and incubated at each organism’s respective temp for 24 h. At the end of incubation, 1 mL of the resulting culture was used for estimation of cell density and pigment quantification, and remaining was used for assaying virulence of the bacterial culture *in vivo* in *C. elegans* model, as described in the preceding section.

**Table 2:** Temperature measurements before and after MW treatment of bacterial suspension

| Duration of MW exposure | *P. aeruginosa* |  | *S. aureus* |  |
| --- | --- | --- | --- | --- |
|  | Before MW treatment (ice treatment for 5 min) | After MW treatment | Before MW treatment (ice treatment for 5 min) | After MW treatment |
| 2 min | 14 | 15.6 | 13.9 | 15.5 |
| 4 min | 14.6 | 15.4 | 14.4 | 15.3 |
| 6 min | 15 | 15.2 | 15.1 | 15.2 |

Immediately after suspending bacterial cells in normal saline, and OD standardization, temp of the bacterial suspensions was found to be 29.5±0.1°C.

### Statistical analysis

All experiments were performed in triplicate, and measurements are reported as mean ± standard deviation (SD) of ≥2 such independent experiments. Statistical significance of the data was evaluated by applying *t*-test using Microsoft Excel^^®^^. *P* values ≤0.05 were considered to be statistically significant.

## Results and Discussion

### Effect of sonic stimulation on bacterial virulence

When *C. elegans* infected with *S. marcescens* were subjected to sonic stimulation (400 Hz), it registered 42% higher survival, as compared to the infected *C. elegans* receiving no sound treatment (Figure 2). Such notable survival benefit was not observed in case of infections caused by remaining two gram-negative pathogens, considering the number of surviving worms on last day of the experiment. However, statistically significant survival benefit was observed in case of *P. aeruginosa* infection till second day (Figure 3), and in case of *C. violaceum* till fourth day (Figure 4). Sonic stimulation could confer 23% better survival on *C. elegans* facing the *S. aureus* challenge (Figure 5). These *in vivo* observations demonstrating effect of sonic stimulation on these bacteria corroborates well with our *in vitro* observations reported previously (Kothari *et al*., 2016) wherein we had shown 400 Hz sound (at 86 dB) to be capable of modulating production of the prodigiosin and violacein pigments in their respective producers. Both these pigments have been reported to possess a variety of biological activities (Liu and Nizet, 2009; Lapenda *et al*., 2015; Hosokawa *et al*., 2016), and their production being under control of QS, sound-induced altered production of these pigments may be taken as an indication of QS machinery of the target bacteria to get modulated under influence of external acoustic field. It should be noted that QS is being actively pursued as a potential target for novel anti-pathogenic therapeutics (Holm and Vikstrom, 2014). Modulation of pigment production in these bacteria may not be solely due to effect of sound on QS machinery. Violacein has been reported not to be solely regulated by QS in *C. violaceum* (de Vasconcelos *et al*., 2003). Altered prodigiosin production might be linked to disturbance of the energy status of *S. marcescens* cells. Haddix *et al*., (2008) suggested a negative regulatory role for prodigiosin in cell and energy production under aerobic growth conditions, and that prodigiosin reduces ATP production by a process termed energy spilling. This process may protect the cell by limiting production of reactive oxygen compounds. Prodigiosin may also possibly act as a mediator of cell death at population stationary phase. Violacein is probably toxic to *C. elegans* (Choi *et al*., 2015), and prodigiosin is also known to exert cytotoxicity (Francisco *et al*., 2007; Dalili *et al*., 2012).

**Figure 2.**
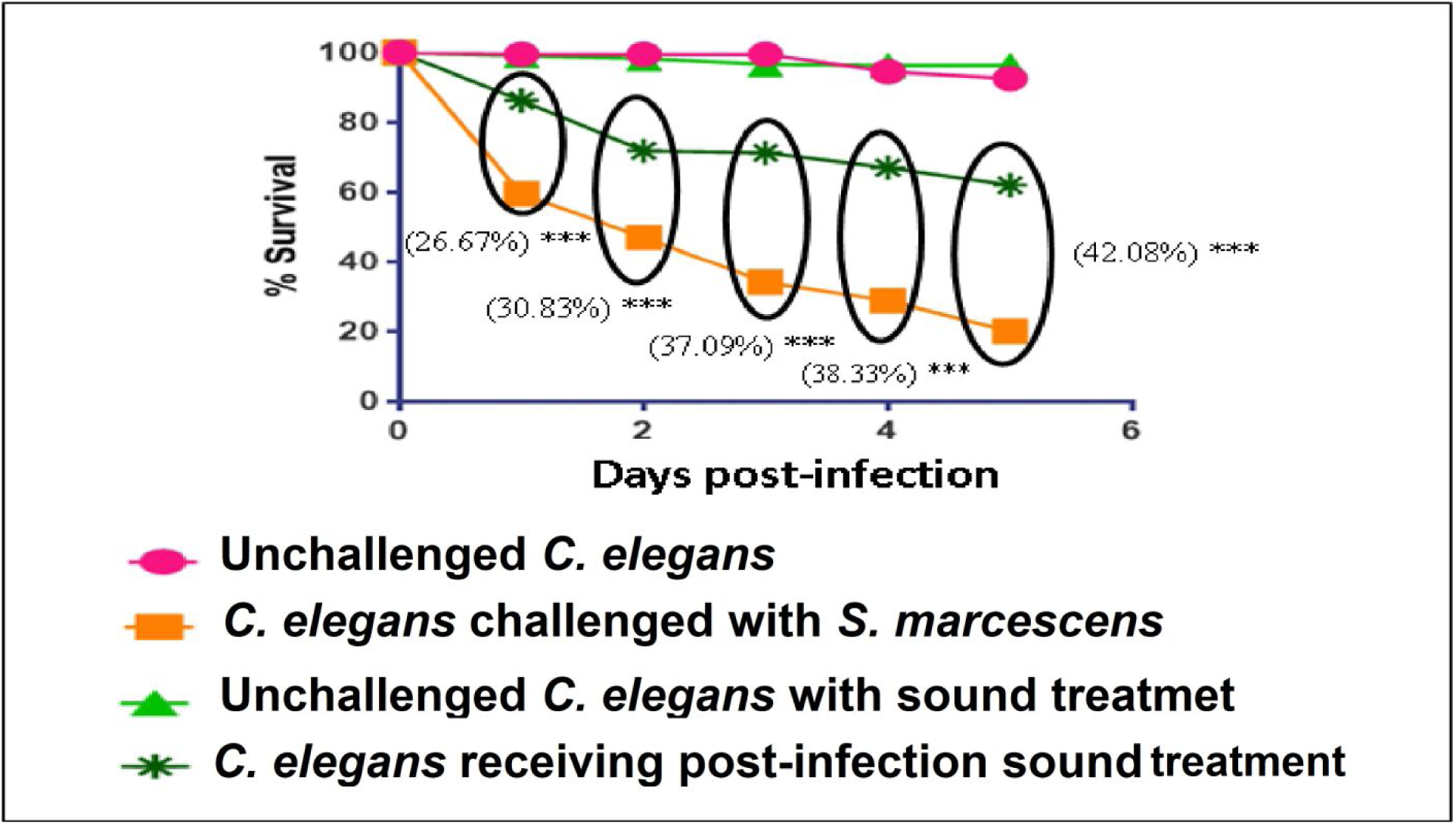
Post-infection sound treatment (400 Hz; 85.5 dB) on *C. elegans* challenged with *S.marcescens*. Values reported are means of 6 independent experiments; ***p<0.001

**Figure 3.**
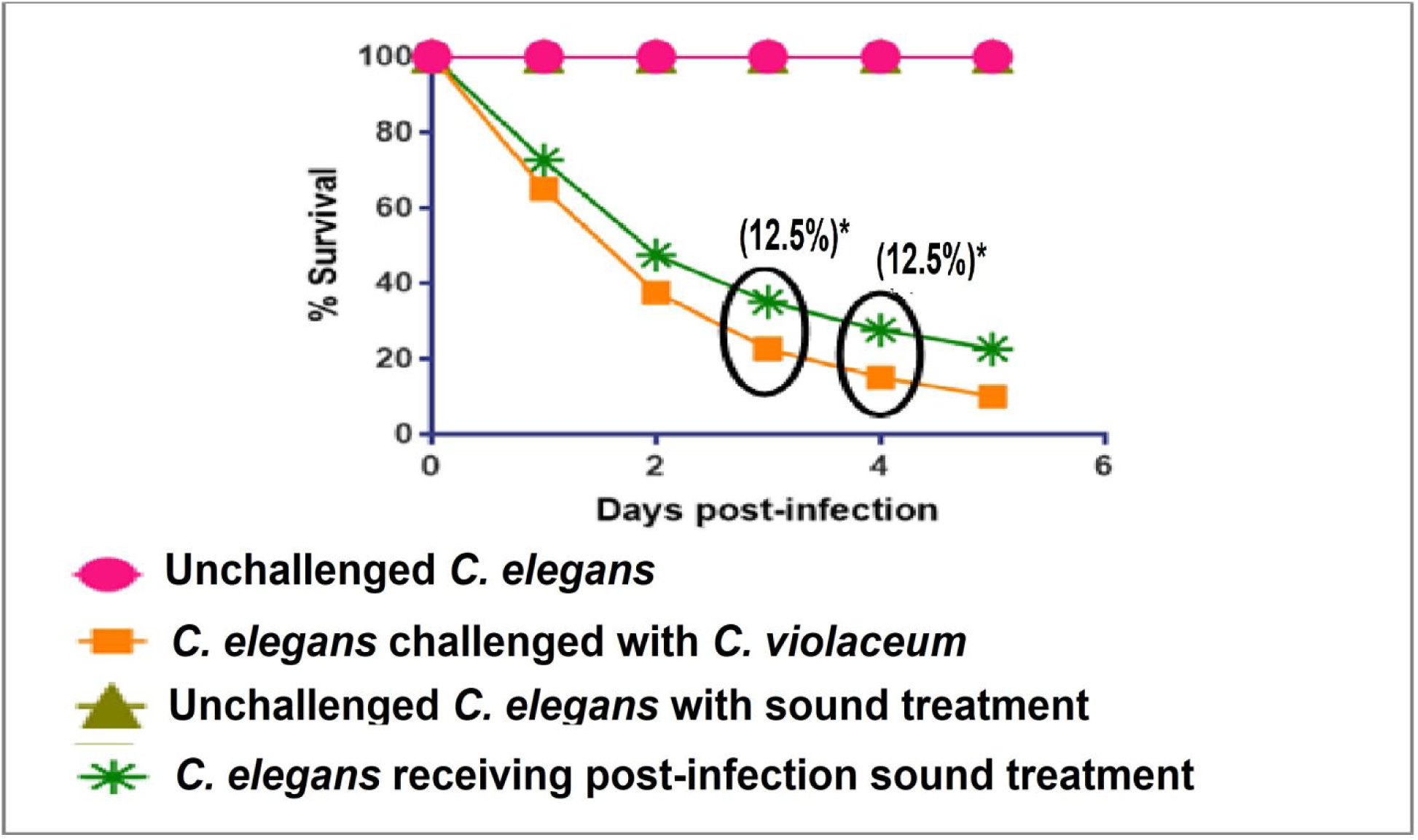
Post-infection sound treatment (400 Hz; 85.5 dB) on *C. elegans* challenged with *P. aeruginosa*. Values reported are means of 2 independent experiments; **p<0.01, ***p<0.001

**Figure 4.**
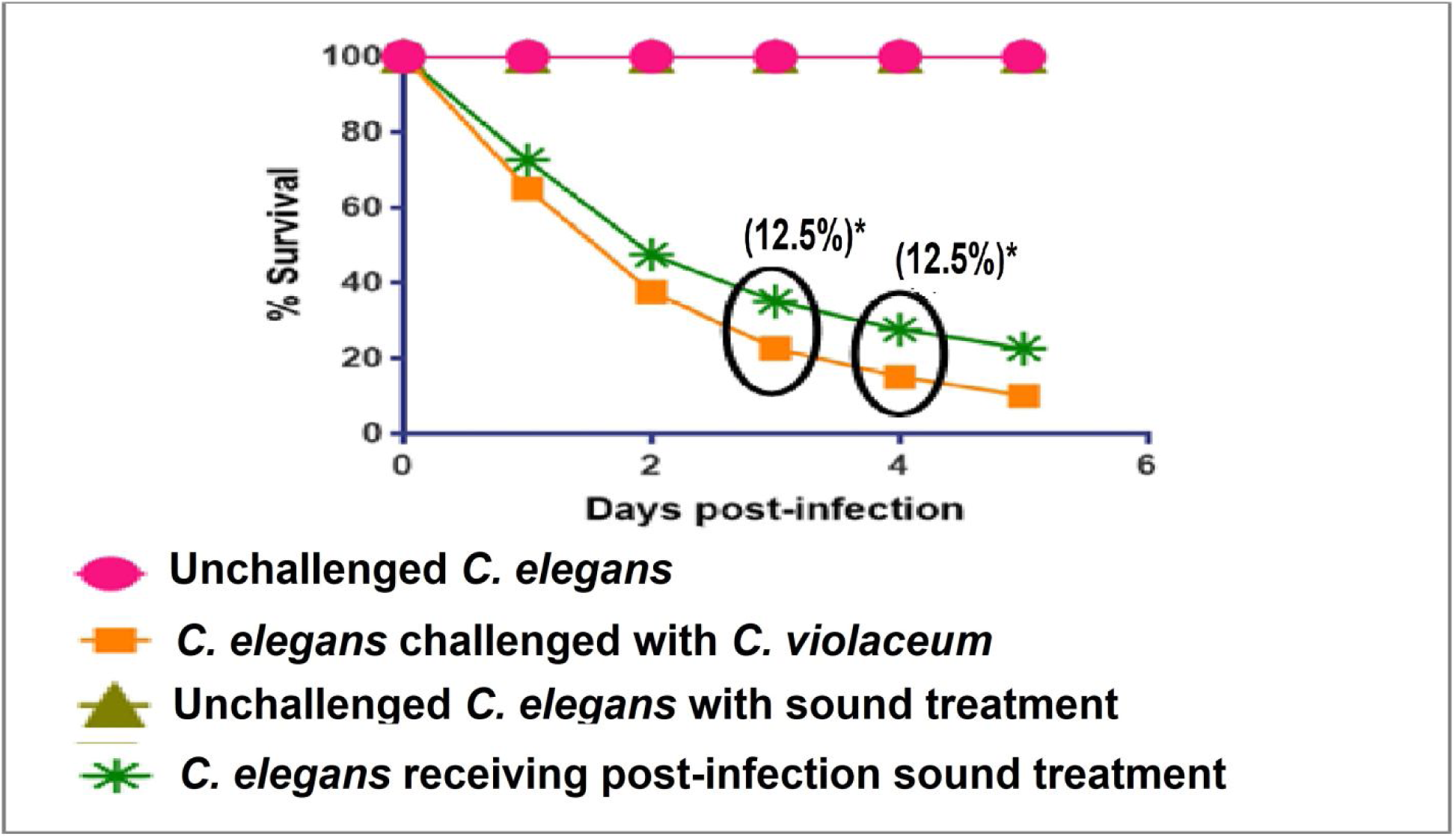
Post-infection sound treatment (400 Hz; 85.5dB) on *C. elegans* challenged with *C. violaceum*. Values reported are means of 2 independent experiments; *p<0.05

**Figure 5.**
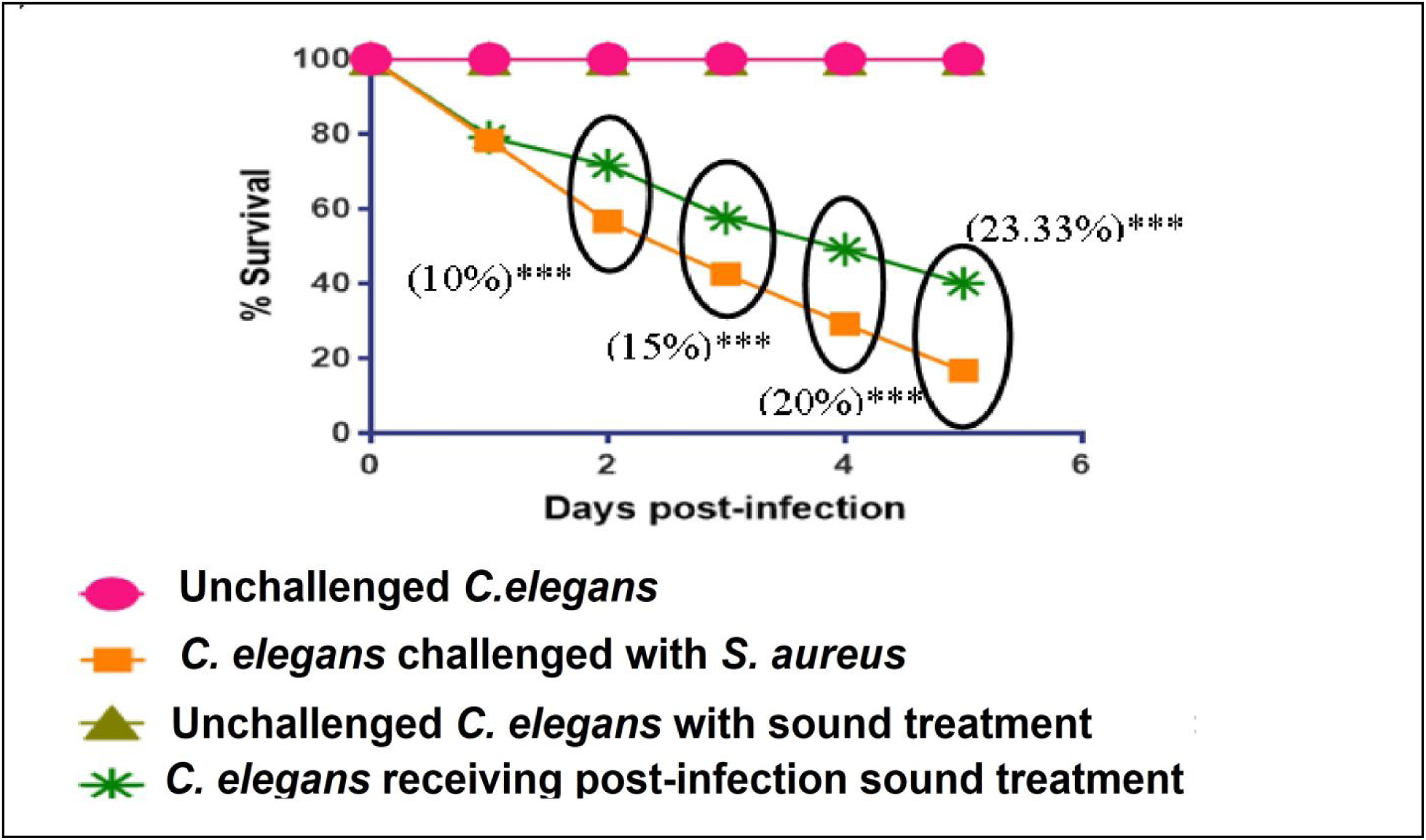
Post-infection sound treatment (400 Hz; 85.5 dB) on *C. elegans* challenged with *S. aureus*. Values reported are means of 3 independent experiments; ***p<0.001

When *C. elegans* infected with different bacteria were exposed to ‘om’ sonic stimulation (Figure 6-9), it registered better survival in face of *S. marcescens* and *S. aureus* challenge, whereas marginal survival benefit seemed to be conferred on this worm owing to sonic stimulation in face of *P. aeruginosa* challenge. In case of *C. violaceum* infection, only a minor (albeit statistically significant) benefit of 7% could be observed on fourth day post-infection.

**Figure 6.**
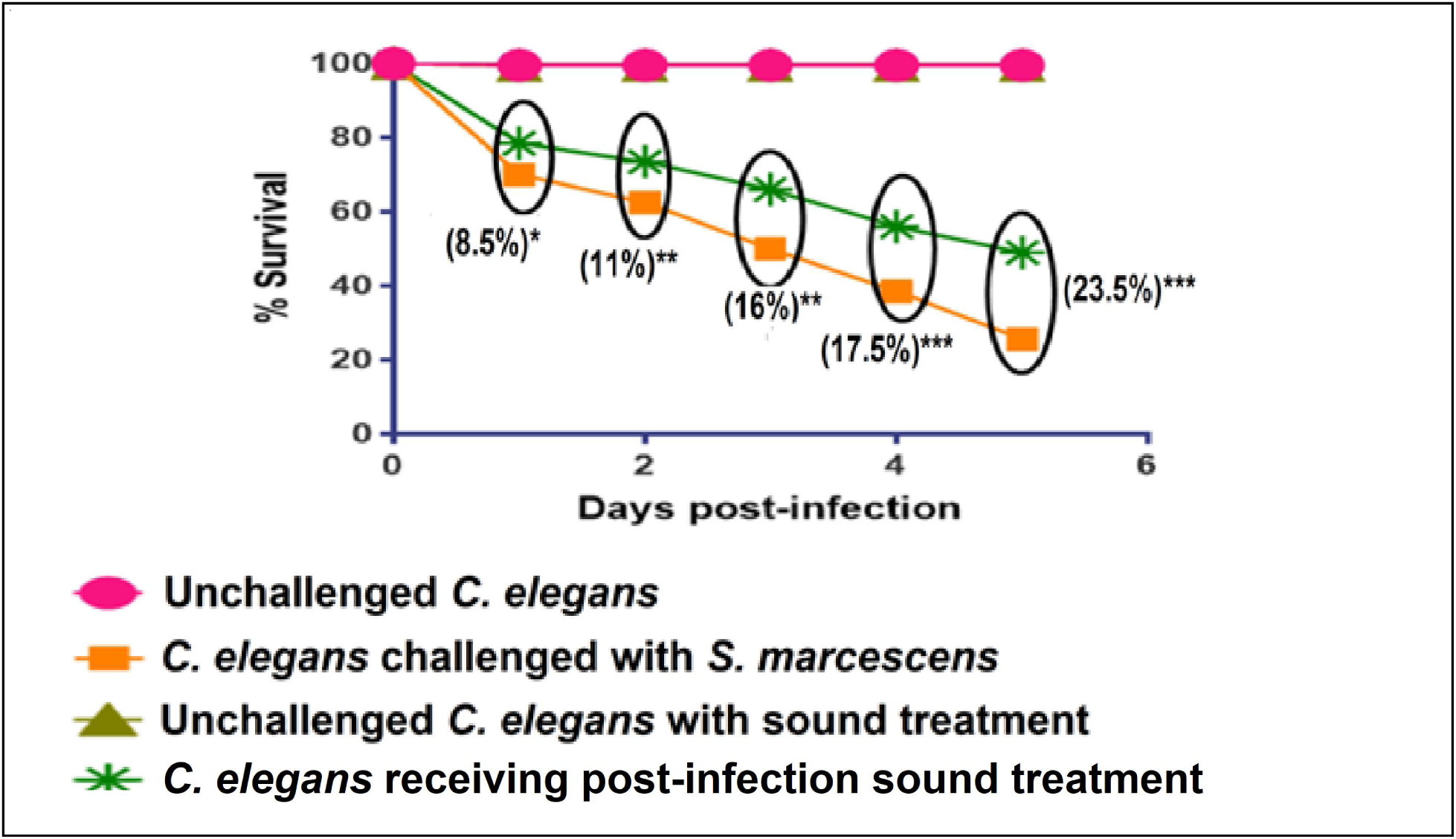
Post-infection sound treatment (Om; 72-75 dB) on *C. elegans* challenged with *S. marcescens*. Values reported are means of 5 independent experiments; *p<0.05, **p<0.01, ***p<0.001

**Figure 7.**
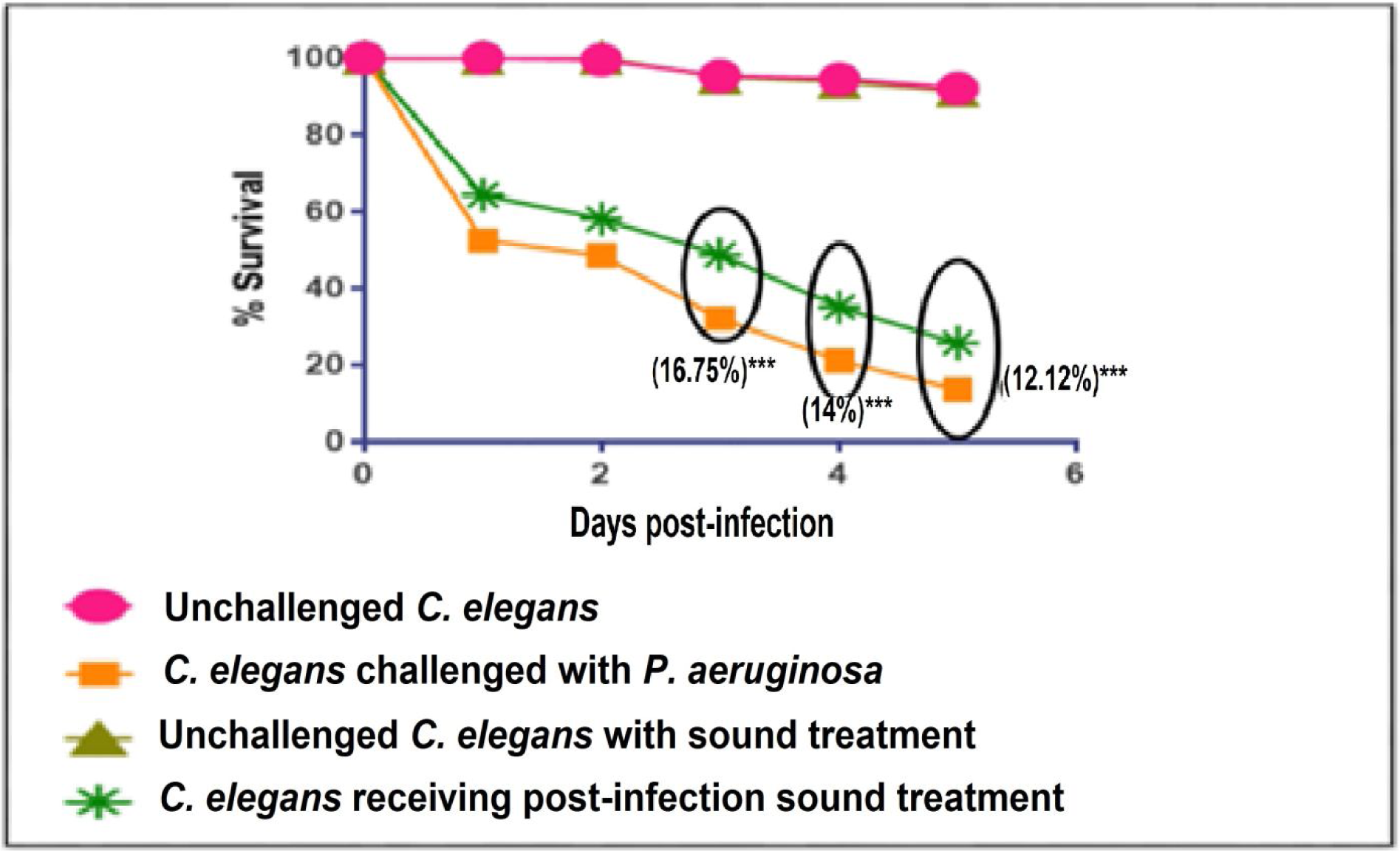
Post-infection sound treatment (Om; 72-75 dB) on *C. elegans* challenged with *P. aeruginosa*. Values reported are means of 6 independent experiments; ***p<0.001

**Figure 8.**
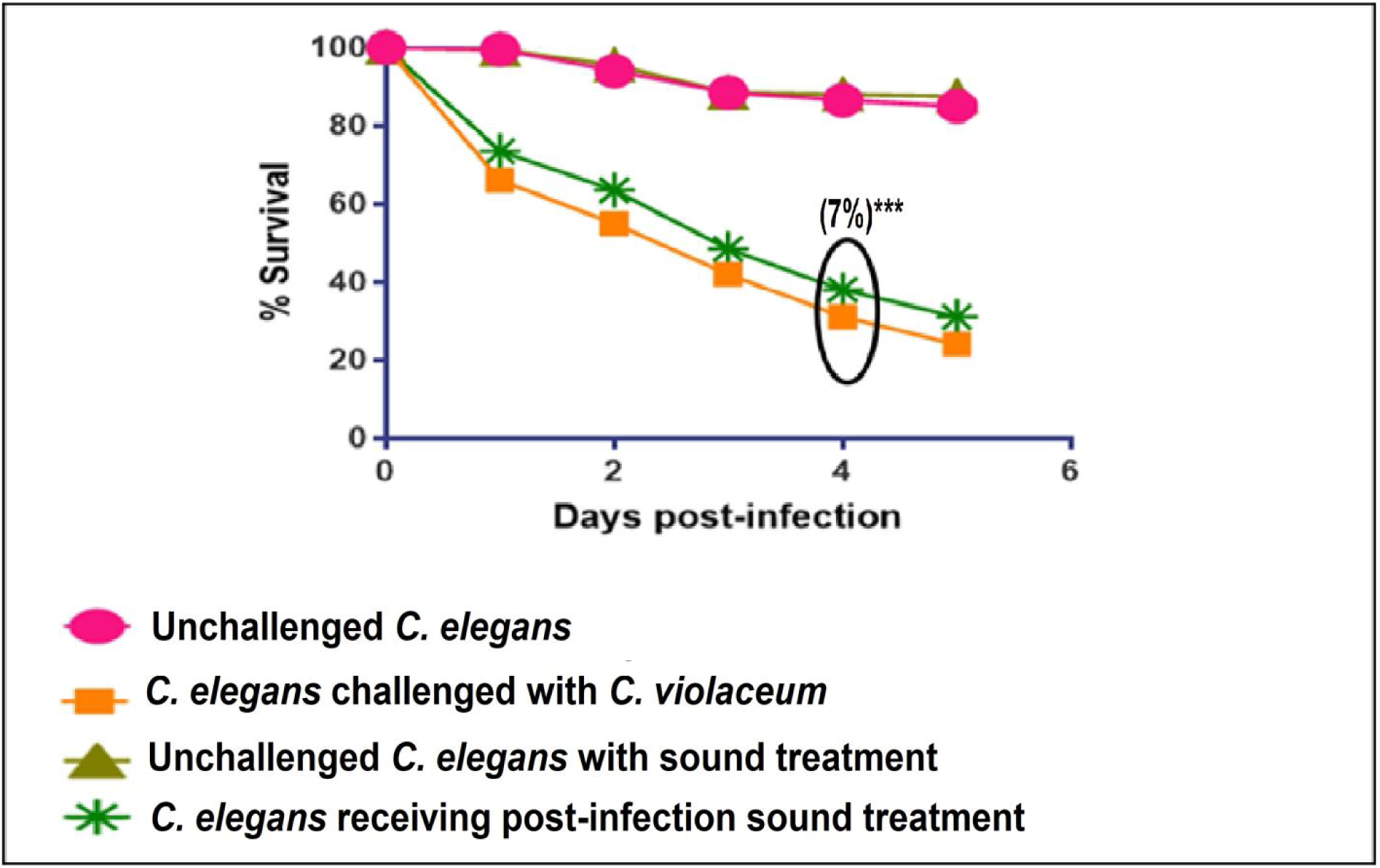
Post-infection sound treatment (Om; 72-75 dB) on *C. elegans* challenged with *C. violaceum*. Values reported are means of 5 independent experiments; ***p<0.001

**Figure 9.**
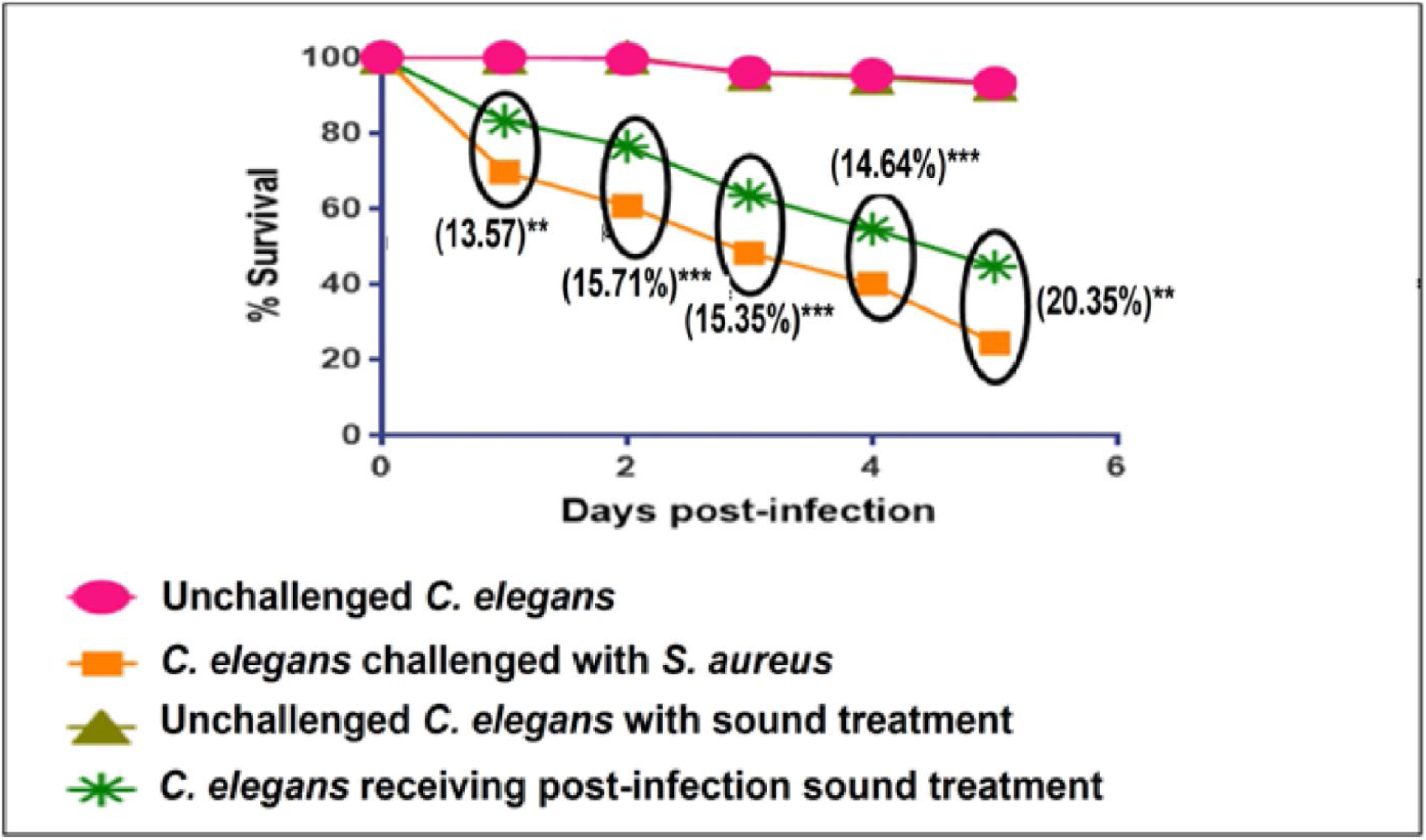
Post-infection sound treatment (Om; 72-75 dB) on *C. elegans* challenged with *S. aureus*. Values reported are means of 7 independent experiments; **p<0.01, ***p<0.001

From results of both types of sounds used to treat *C. elegans* post-bacterial infection, it seems that infection caused by *S. marcescens* and *S. aureus* respond better to the sonic therapy, whereas outcome of *C. violaceum* infection does not seem to get affected much due to sonic stimulation. In any of these experiments, sonic treatment was not found to have any positive or negative effect on overall survival percentage of *C. elegans*, in absence of bacterial challenge, suggesting that all the observed effect is likely be due to influence of sound on the bacterial pathogen. This is to say that the improved survival of the nematode worm experiencing sonotherapy, in face of bacterial challenge, is very much likely to be due to sound-responsiveness of the bacteria. Insensitivity of wild type *C. elegans* to low-pressure ultrasound has been reported by Ibsen *et al*. (2015).

This property of different bacteria to respond to sonic frequencies has been reported earlier. First such report (Matsuhashi *et al*., 1998) described not only the ability of *Bacillus carboniphilus* to respond to sound, but also the ability of *B. subtilis* to produce sound waves. Gu *et al*. (2013) described alteration in growth and physiological characters of *E. coli* upon exposure to external sound field. Later they (Gu *et al*., 2017) studied differential gene expression in *E.coli* K12 exposed to sound. First report describing effect of sonic stimulation on bacteria at the whole transcriptome level was made public in Jan 2017 by our group (https://www.biorxiv.org/content/early/2017/01/04/098186). Liu *et al*. (2016) described germination-promoting effect of sonic stimulation (5 kHz at 90 dB) on *Bacillus* endospores. Though now it is known that bacteria and other microbes respond to external sound exposure, which is described in terms of alteration in their growth, metabolism, gene expression etc., the precise mechanism governing this phenomenon of sound-responsiveness among bacteria remains to be found out. The acoustic energy absorbed by bacteria may be postulated to modulate their membrane permeability and enzyme activities. Liu *et al* (2016) indicated the possibility of increased release of chemical mediators of QS in sound-exposed endospores. We have also shown altered production of QS-regulated pigments in sound-exposed bacterial cultures (Kothari *et al*., 2017; Kothari *et al*., 2018). QS being an important regulator of virulence (Natrah *et al*., 2011), any chemical or physical agents capable of modulating QS can be expected to modulate bacterial virulence too. Sound waves may exert their effect on bacteria by modulation of the chemotaxis (Gu *et al*., 2017) and/or glucose metabolism pathway (Joshi *et al*., 2018). These effects may be believed to be dependent on factors like size/ shape of the test bacteria, their growth phase (e.g. logarithmic vs. stationary), etc. Aspect ratio of the bacterial cells/ spores may play an important role in determining the magnitude of specific absorption rate toward sound waves (Liu *et al*., 2016).

### Effect of sonic stimulation on *P. melaninogenica* and *B. bifidum*

First we exposed *P. melaninogenica* to ‘Om’ sound at two different intensities i.e. 70-75 dB and 80-85 dB. This sound at 70-75 dB promoted growth of this bacterium by 5.88-62.93%. However this growth-promoting effect was observed 4 times out of 6 times we did this experiment (Table 3). These results can be said to be inconclusive, as outcome of all the six experiments did not match, and among those four experiments, when growth of the sonic stimulated culture was found to be statistically significantly higher, the range of magnitude of effect (expressed in %) was a wide range. At present, we are not able to understand and explain the reasons behind these inconclusive results. The same ‘Om’ sound at bit higher intensity of 80-85 dB failed to exert any influence on this bacterium’s growth. Based on these results, we selected this sound at 70-75 dB for treating *B. bifidum*, but no effect was observed.

**Table 3.** Effect of ‘om’ on *P. melaninogenica* and *B. bifidum*

| Sr. no. | Organism | Sound intensity (dB) | Experiment No. | Control (OD<sub>764</sub>) (Mean ± SD) | Experiment (OD<sub>764</sub>) (Mean ± SD) | % change (Mean ± SD) |
| --- | --- | --- | --- | --- | --- | --- |
| 1 | *P. melaninogenica* | 70-75 | 1 | 0.56 ± 0.005 | 0.64±0.01 | 14.13**±3.66 |
|  |  |  | 2 | 0.76 ± 0.02 | 0.83±0.02 | 11.37*±4.92 |
|  |  |  | 3 | 0.59 ± 0.007 | 0.63±0 | 5.88*±1.25 |
|  |  |  | 4 | 0.65 ± 0.04 | 0.66±0.12 | 1.11±12.98 |
|  |  |  | 5 | 0.53 ± 0.05 | 0.54±0.02 | 3.63±15.06 |
|  |  |  | 6 | 0.58 ± 0.014 | 0.90±0.02 | 62.93**±0.31 |
|  |  | 80-85 | 1 | 0.65 ± 0.08 | 0.65±0.07 | 0.63±20.72 |
|  |  |  | 2 | 0.64 ± 0.07 | 0.56 ± 0.10 | -10.88 ± 28.24 |
| 2 | *B. bifidum* | 71-75 | 1 | 0.72±0.007 | 0.72±0.01 | -0.67±2.91 |
|  |  |  | 2 | 0.73±0.03 | 0.85±0.01 | 15.82*±7.49 |
|  |  |  | 3 | 0.3±0.04 | 0.40±0.02 | 35.35±28.56 |

\*p<0.05, \*\*p<0.01; '-' sign indicates a decrease over control

Since the ‘Om’ sound was able to promote growth of *P. melaninogenica* 4 out of 6 times, we identified the major frequency components of this sound, by performing a temporal frequency analysis (TFFT) using NCH WavePad v 7.13. This analysis resulted in identification of five major frequency bands in the poly frequency sound ‘om’ viz. 136.1 Hz, 270 Hz, 405 Hz, 540 Hz, and 810 Hz. Following this we generated separate sound beeps for each of these frequencies, and a file containing their combination. These all were pulsed sound files, with a time gap of 1 second between two consecutive beeps. Effect of these six sound types was also checked on these two bacteria, and in none of the case, any reproducible effect on their growth could be observed (data not shown).

### Effect of MW treatment on bacterial virulence

We also investigated effect of low power MW radiation on two of the most notorious human-pathogenic bacteria i.e. *P. aeruginosa*, and *S. aureus*. When MW-treated inoculum of these bacteria was inoculated into their respective growth media, at the end of incubation period, growth resulting from the MW-exposed inoculum was not found to be different from that resulting from MW-unexposed inoculum; similarly MW treatment was not found to alter pigment production in these bacteria (Figure 10 and Figure 12). However, virulence of *S. aureus* culture subjected to 2 min MW exposure was significantly found to be reduced in *in vivo* assay employing the nematode host, wherein the MW-treated *S. aureus* culture could kill lesser number of worms till fourth day, as compared to the control bacterial culture (Figure 11). Virulence of *P. aeruginosa* was not affected by MW treatment (Figure 13).

**Figure 10.**
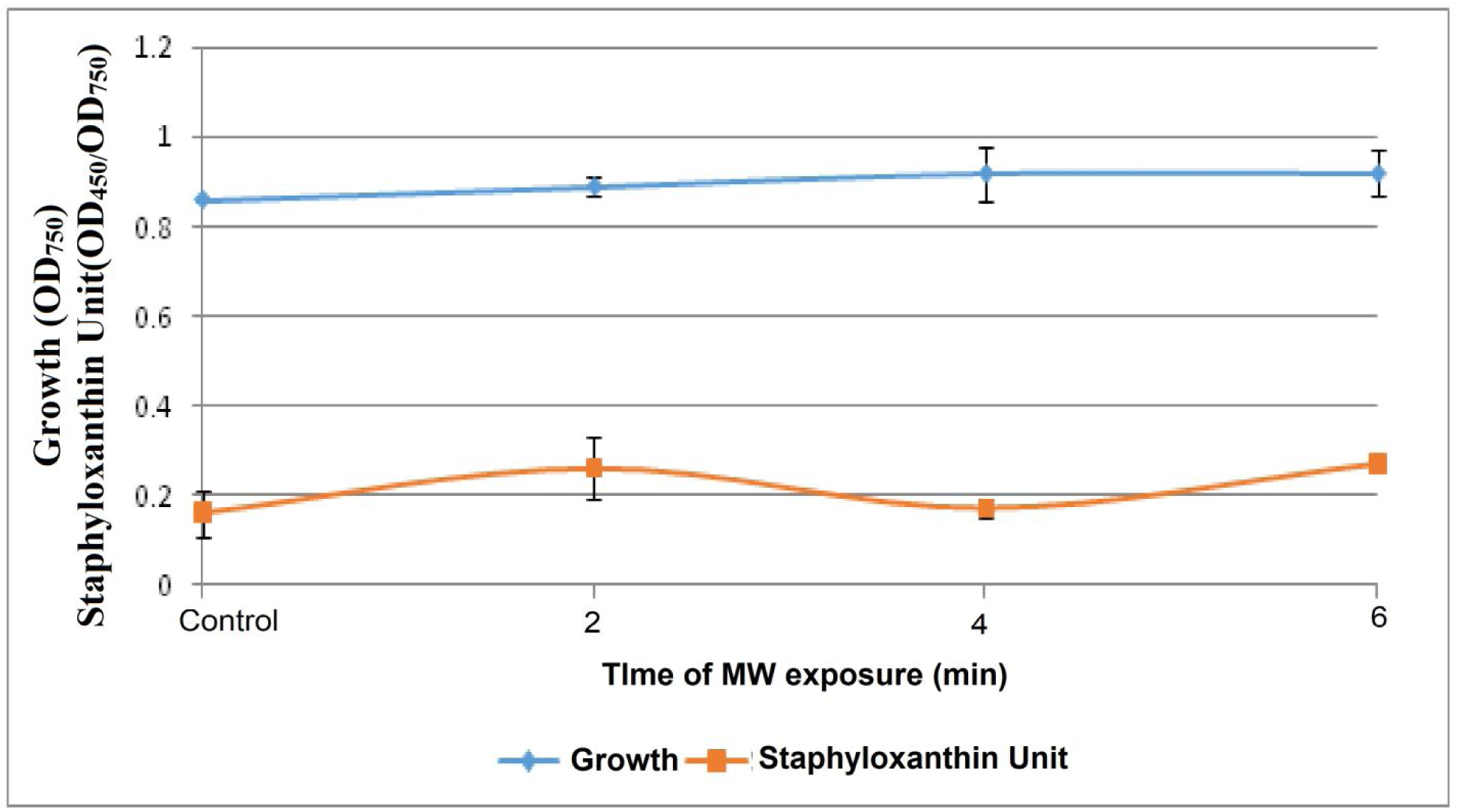
*S. aureus* growth and pigment production were not affected by MW treatment. Bacterial growth was measured as OD_764_; OD of staphyloxanthin was measured at 450 nm, and Staphyloxanthin Unit was calculated as the ratio OD_450_/OD_750_ (indication of staphyloxanthin production per unit of growth). Pigment (staphyloxanthin) extraction and quantification was done as described in Song *et al*. (2009). Bacterial culture not exposed to MW served as control.

**Figure 11:**
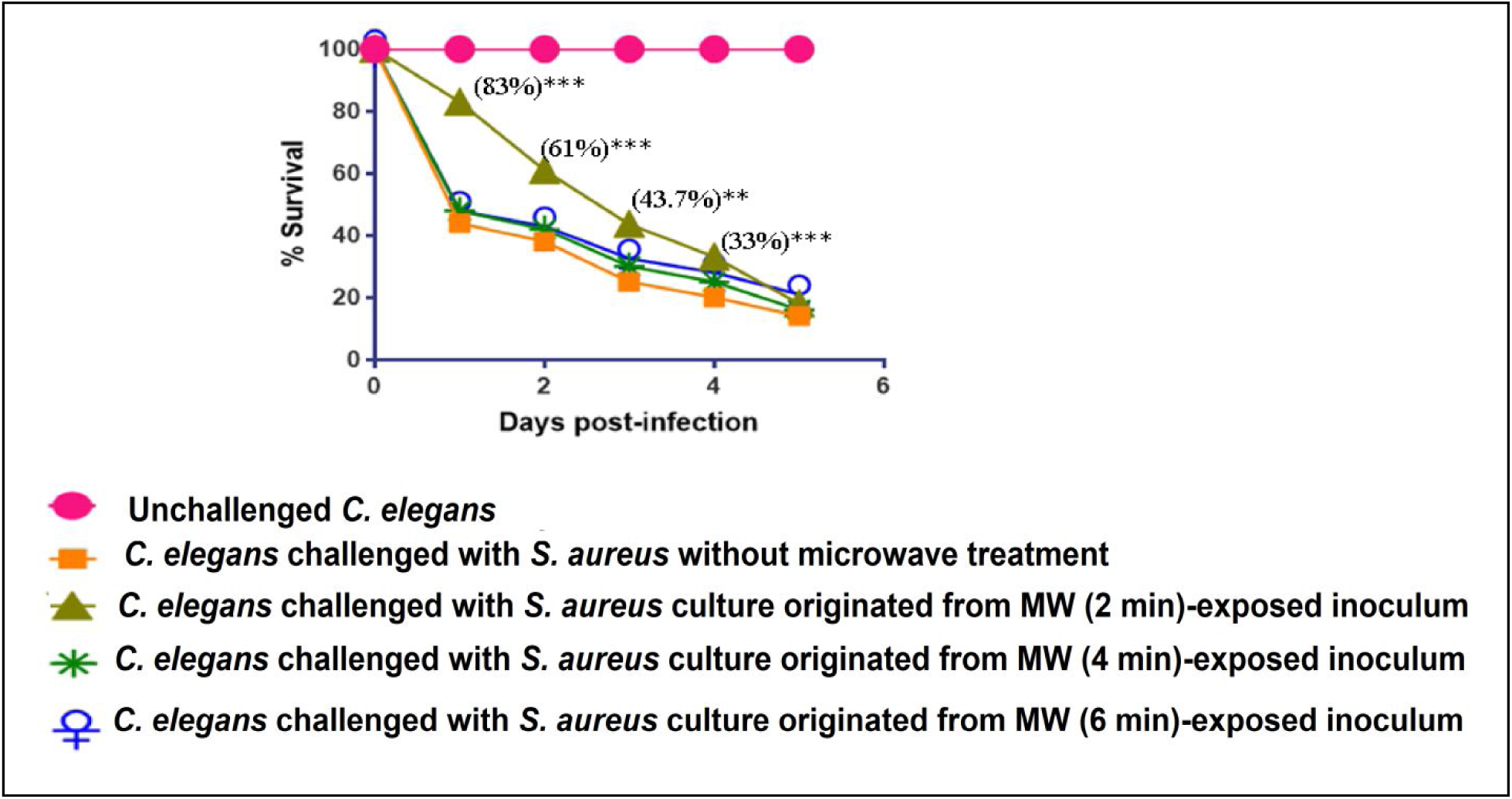
*S. aureus* culture resulting from MW (2 min)-exposed inoculum expressed reduced virulence towards *C. elegans*. Values reported are means of 4 independent experiments; **p<0.01, ***p<0.001

**Figure 12.**
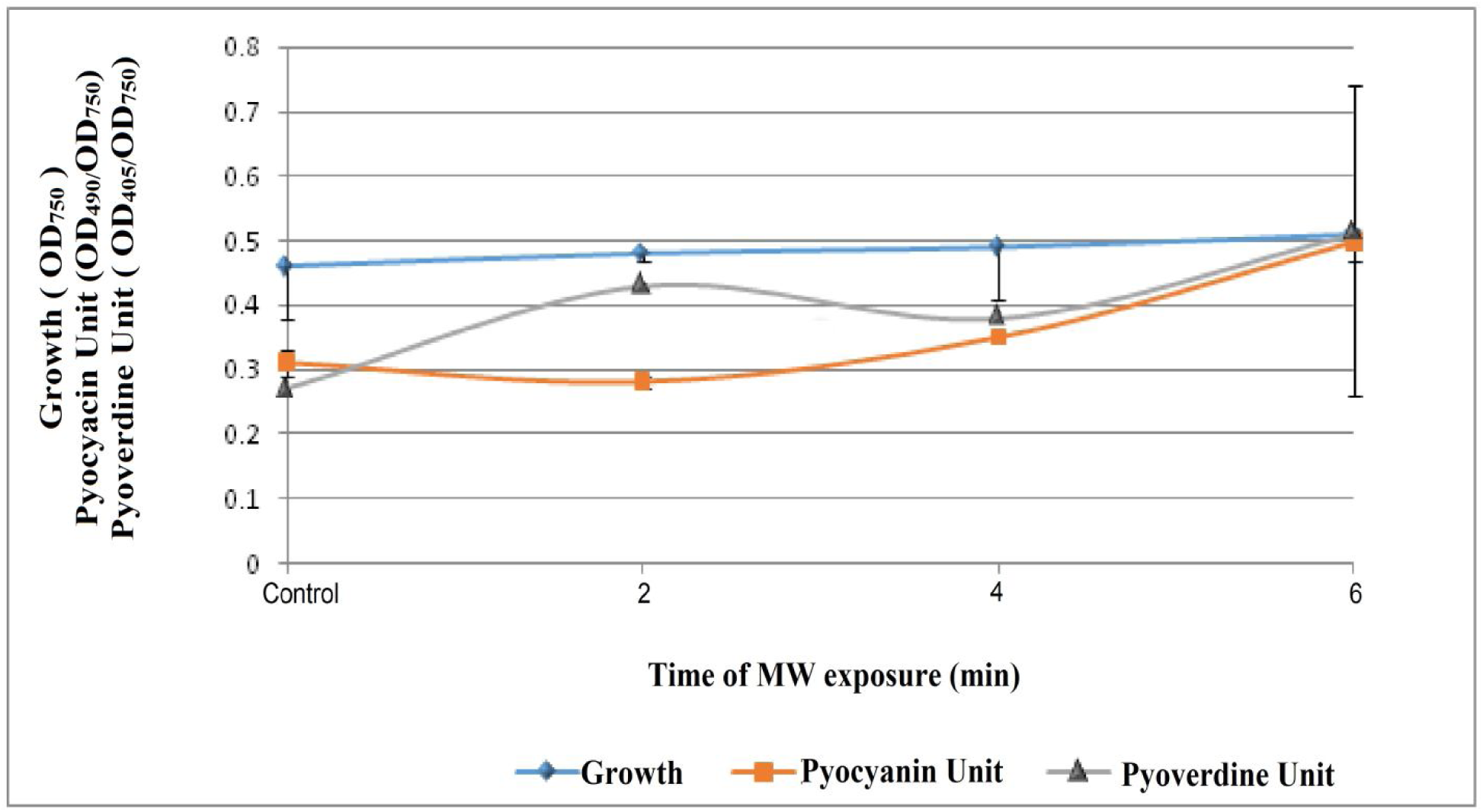
MW treatment did not affect *P. aeruginosa* growth and pigment production. Bacterial growth was measured as OD_750_; OD of pyoverdine was measured at 405 nm, pyocyanin was measured at 520 nm. Pyoverdine Unit was calculated as the ratio OD_405_/OD_750_ (an indication of pyoverdine production per unit of growth); Pyocyanin Unit was calculated as the ratio OD_490_/OD_750_ (an indication of pyocyanin production per unit of growth); Bacterial culture receiving no MW exposure served as acontrol. Extraction and quantification of the pigments pyoverdine and pyocyanin was carried out as described in El-Fouly *et al*. (2015) and Unni *et al*. (2014) respectively.

**Figure 13.**
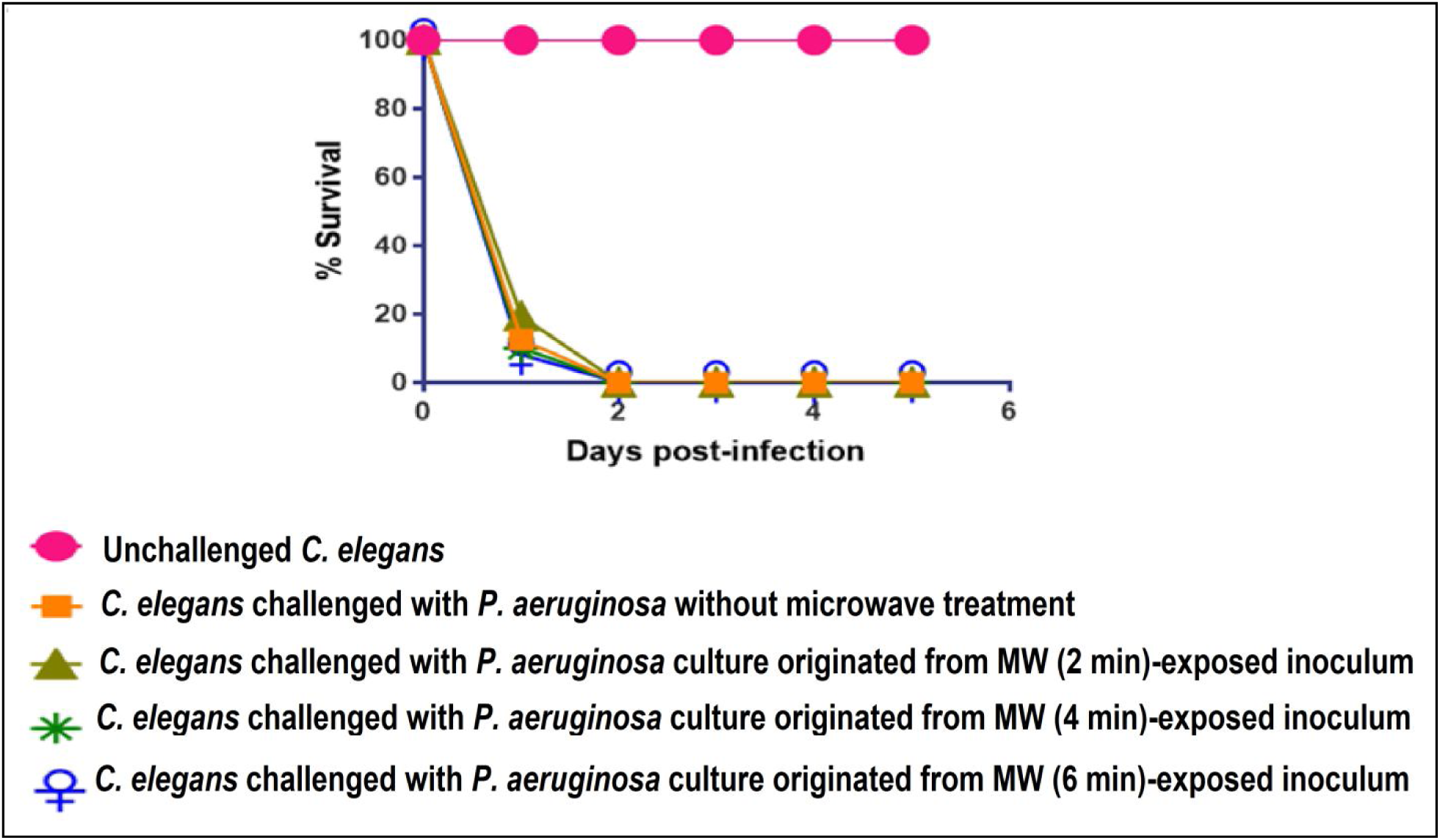
Microwave treatment was not found to alter virulence of *P. aeruginosa* towards *C. elegans*. Values reported are means of 4 independent experiments.

In the present study, assessment of bacterial virulence towards *C. elegans* was made using the bacterial cell population originated from MW treated inoculum, and not directly on the MW treated cells. Therefore, the alterations in virulence might be believed to be transferred from the originally MW treated cells to their daughter cells (who did not receive direct MW exposure). As in this study, thermal effect of MW was avoided by putting the inoculum in ice during MW treatment, whatever alterations have been observed (i.e. in case of 2-min MW exposure to *S. aureus*) are most likely a result of MW specific effects (athermal effects), as these changes occurred in absence of any noticeable heat generation. Exact mechanism by which MW exert their athermal effects on biological systems is still not clear.

Despite the fact that biological significance of MW-specific athermal effects has not been established beyond doubt, reports indicating their possible existence (Dreyfuss and Chipley 1980; Copty *et al*., 2006; Carta and Desogus, 2010) has attracted considerable attention. As low intensity MW is believed not to possess sufficient energy for breaking chemical bonds directly, alternative mechanisms of interaction between MW and biological entities are likely to prevail. This situation is somewhat similar to the status of use of ‘cold plasma’ for germicidal purpose, wherein mode of action is not very clear, that how inactivation of bacteria is achieved by non-thermal plasmas (Morent and De Geyter, 2011; Tipa *et al*., 2011).

This study attempted to investigate the effect of sonic stimulation, and low-power MW radiation on bacterial virulence. Both these treatments (at least with one test bacterium) were found to be capable of modulating bacterial virulence towards the nematode worm *C. elegans*, more so with the sonic approach. However not enough research has been reported regarding biological effects of audible range of sound, as well as, low-power MW, particularly at the cellular level. Microorganisms owing to their ease of handling in lab, rapid growth rate, etc. offer a good model for investigation in this area. Though reports describing effect of sound waves or MW on different microbes are there in literature, how they exert their effect remains obscure. Further these being physical agents (like UV), their effects may follow a random pattern, and reproducible results may be difficult to obtain (as happened during our experiments with *Prevotella melaninogenica)*. Further, many of their effects may be reversible, for example a sonic stimulated culture may revert back to the *normal* behavior, soon after removal of the external acoustic field. Reversible nature of MW-induced mutations has previously been indicated by us (Raval *et al*., 2014). Though underdeveloped at present, further systematic research in this area may pave the way for development of novel therapeutic applications. Some reports do indicate therapeutic utility of acoustic waves (i.e. sonotherapy), and MW, their use for management of microbial infections has yet not been possible. For example, vibroacoustic therapy is being described as a new sound technology that uses audible sound vibrations to reduce symptoms, invoke relaxation, and alleviate stress. Use of this technology (e.g. in pain management) is based on the fact that external vibration can influence body function (Boyd-Brewer and McCaffrey, 2004). Music has also been proposed as a therapy for anxiety and pain (Negrete, 2011).However, use of acoustic vibrations for microbiological purposes will be possible only after this area of knowledge gains sufficient attention from the research community, and mechanistic details of how microbes and sound waves interact are understood. Our understanding of whether and how non-auditory eukaryotic cells are affected by sonic stimuli is still fragmentary. Lestard *et al*. (2013) suggested that music can alter morpho-functional parameters, such as cell size and granularity in cultured cells, and that music can interfere with hormone binding to their targets, indicating that music (audible sounds) could modulate physiological and pathophysiological processes. Our understanding of how non-auditory cells are affected by sonic stimuli may enable the scientific community to devise vibroacoustic therapy for management of microbial infections, and also for disease conditions which are shown to have correlation with alterations in human microbiota.

Though sonic stimulation in this study was not found to alter growth of *P. melaninogenica* and *B. bifidum*, it did attenuate virulence of *S. marcescens* towards *C. elegans. S. marcescens* is a member of the family Enterobacteriaceae, whose abundance has been reported to be positively associated with the severity of postural instability and gait difficulty in PD patients. *C. elegans* is a small soil worm or nematode, and it shares a common ancestor with humans that lived in the pre-Cambrian era (500-600 million years ago). This ancestor is referred to as the urbilaterian ancestor, as is believed to be the relative of all bilaterally symmetric, multicellular organisms on the earth, including invertebrates and vertebrates (http://www.people.ku.edu/~erikl/Lundquist_Lab/Why_study_C._elegans.html). It also shares some 40% similarity with human genome. Our experiments showed sonic treatment to confer notable benefit on *C. elegans* upon challenge with *S. marcescens*. Latter has been known to be involved in human wound infections on skin, and *C. elegans* epidermis has been proposed as a model skin (Chisholm and Xu, 2012).

To the best of our awareness, this is the first report, describing sonic/ MW exposure to cause alteration in bacterial virulence towards the nematode *C. elegans*. Further experiment to understand the molecular basis of this phenomenon are warranted. A thorough understanding of the mechanism how EMR or sonic stimulation affects bacterial physiology and virulence is required, for development of any useful therapeutic application of such non-invasive approaches to tackle bacterial infections.

